# Mouse frontal cortex nonlinearly encodes sensory, choice and outcome signals

**DOI:** 10.1101/2023.05.11.539851

**Authors:** Lauren E. Wool, Armin Lak, Matteo Carandini, Kenneth D. Harris

## Abstract

Frontal area MOs (secondary motor area) is a key brain structure in rodents for making decisions based on sensory evidence and on reward value. In behavioral tasks, its neurons can encode sensory stimuli, upcoming choices, expected rewards, ongoing actions, and recent outcomes. However, the information encoded, and the nature of the resulting code, may depend on the task being performed. We recorded MOs population activity using two-photon calcium imaging, in a task requiring mice to integrate sensory evidence with reward value. Mice turned a wheel to report the location of a visual stimulus following a delay period, to receive a reward whose size varied over trial blocks. MOs neurons encoded multiple task variables, but not all of those seen in other tasks. In the delay period, the MOs population strongly encoded the stimulus side but did not significantly encode the reward-size block. A correlation of MOs activity with upcoming choice could be explained by a common effect of stimulus on those two correlates. After the wheel turn and the feedback, the MOs population encoded choice side and choice outcome jointly and nonlinearly according to an exclusive-or (XOR) operation. This nonlinear operation would allow a downstream linear decoder to infer the correct choice side (i.e., the side that would have been rewarded) even on zero contrast trials, when there had been no visible stimulus. These results indicate that MOs neurons flexibly encode some but not all variables that determine behavior, depending on task. Moreover, they reveal that MOs activity can reflect a nonlinear combination of these behavioral variables, allowing simple linear inference of task events that would not have been directly observable.

## Introduction

The secondary motor area (MOs) in mice is a key frontal cortical structure for making decisions based on sensory evidence and reward (Barthas and Kwan, 2017). Inactivation of MOs affects performance in multiple tasks, involving multiple sensory modalities and behavioral outputs (Erlich et al., 2011, 2015; Sul et al., 2011; Guo et al., 2014; Hanks et al., 2015; Goard et al., 2016; Coen et al., 2021; Kondo and Matsuzaki, 2021; Zatka-Haas et al., 2021; Atilgan et al., 2022). Recordings in MOs have shown diverse correlates of task variables including sensory inputs, rewards, and choices. Choices can often be predicted from MOs activity earlier than from activity in other brain regions (Erlich et al., 2011; Sul et al., 2011; Murakami et al., 2014; Li et al., 2016; Siniscalchi et al., 2016, 2019; Bari et al., 2019; Jiang et al., 2019; Steinmetz et al., 2019; Kondo and Matsuzaki, 2021; Shin et al., 2021), suggesting that MOs plays a key role in decision-making in rodents.

In certain tasks, MOs shows correlates of upcoming choice and expected reward before the relevant action is performed. For instance, in a maze-based two-arm bandit task (Sul et al., 2011), MOs can encode information on which choice will deliver a larger reward. Moreover, in tasks that require memorizing a sensory stimulus, MOs encodes choice during the delay period between stimulus and action (Erlich et al., 2011; Li et al., 2016).

The fact that MOs exhibits correlates of these task features in specific tasks, however, does not necessarily imply its role is always to encode them. To understand the role a particular brain area performs it is necessary to understand how the information it encodes varies depending on the particular task being performed. Furthermore, when task variables correlate with each other, neural correlates of one variable can potentially be explained by common effects of another. For example, a neural correlate of the forthcoming choice during a delay period might reflect encoding of a planned action, but it might also reflect encoding of a sensory stimulus that is used to guide the choice.

A related question is the way in which task variables are encoded in MOs population activity. Do different neurons encode different task variables, or can single neurons multiplex multiple variables to form a mixed representation—and if so, does this occur in a nonlinear manner (Rigotti et al., 2013; Bernardi et al., 2020)? This question of coding linearity is particularly important in tasks where the behavioral variables are themselves nonlinearly related, as it affects the conclusions that researchers can draw from observing the encoding of a variable—and, indeed, the strategies that downstream structures might use to take advantage of MOs signals.

Here we investigate neural activity in MOs in a decision task requiring mice to integrate sensory information with reward value. We find that MOs encodes different task variables than in other tasks, and encodes two of these variables in a nonlinear exclusive-or (XOR) fashion. In our task, mice made choices based on random visual stimuli, for rewards whose size varied in randomly shifting blocks (Lak et al., 2020). Before responding, mice had to wait over a delay period (during which the stimulus remained visible) which was ended by an auditory “Go” cue. This delay enables separate testing for neural correlates of stimulus, choice, and reward, because these occur at different timepoints. Unlike a bandit task (Sul et al., 2011), in our task MOs did not significantly encode the reward block (i.e., information on which side will deliver a larger reward), and significantly encoded available reward size only after the choice was made. Also, unlike in memorization tasks (Erlich et al., 2011; Li et al., 2016), an apparent encoding of upcoming choice during the delay period could be explained by a common effect of the sensory stimulus on choice and neural activity. After wheel turn and feedback onset, the MOs population encoded choice side and choice outcome jointly and nonlinearly according to an exclusive-or (XOR) operation. This nonlinear operation would allow a downstream linear decoder to infer the correct choice side (i.e., the side that would have been rewarded) even on zero contrast trials, when there had been no visible stimulus. These results indicate that MOs neurons flexibly encode some but not all variables that determine behavior, depending on task. Moreover, they reveal that MOs activity can reflect a nonlinear combination of these behavioral variables, allowing simple linear inference of task events that would not have been directly observable.

## Results

### Mouse choices are guided by stimuli and rewards

We recorded neural populations from frontal area MOs in a two-alternative visual decision-making task with unequal rewards (Lak et al., 2020). Mice were head-fixed and surrounded by 3 screens. On each trial, a grating stimulus of randomly varying contrast appeared randomly on the right or the left screen. Mice turned a steering wheel with their forepaws to report the stimulus location for a liquid reward (Burgess et al., 2017) (Figure 1A). In our version of the task, mice had to keep the wheel still for a delay period of 0.8–1.0 s after stimulus onset (Figure 1B). Any wheel movement during this period extended the delay, and these trials were excluded from analysis. The end of the delay period was signaled by an auditory Go cue indicating that mice should turn the wheel to drive the stimulus into the center screen. Mice received a 10% sucrose water reward for correct responses, or an auditory white noise burst and time out on error trials. The reward differed between left and right choices, with one side receiving twice as much liquid (2.4 versus 1.2 µL). The high-reward side switched without warning and without cue in “reward blocks” of 125–225 trials (Figure 1C). Regardless of block, the stimulus always had a 50% chance of appearing on either side, and a reward was only given for choosing the correct stimulus side (Lak et al., 2020).

**Figure 1.**
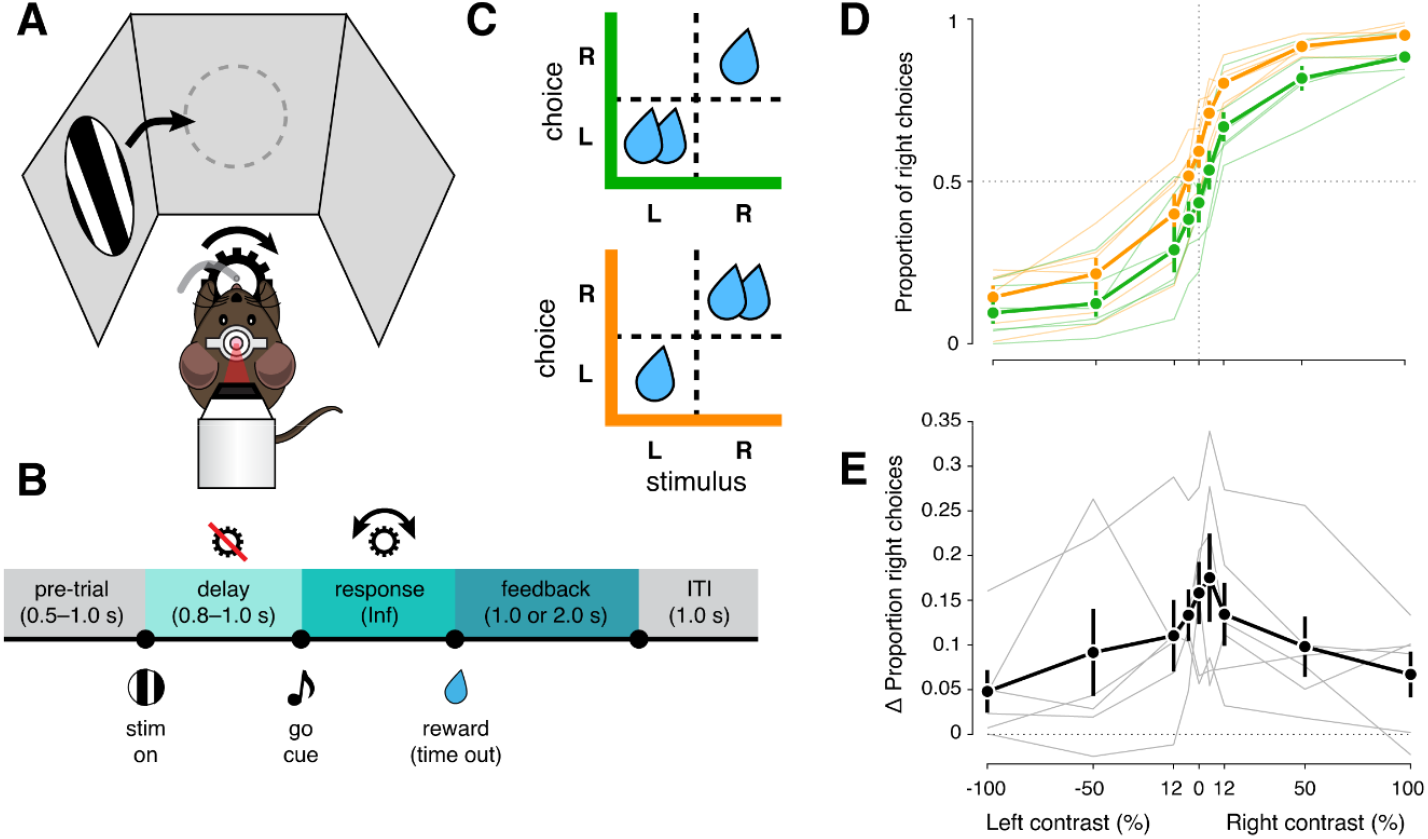
Mouse choices are guided by stimuli and rewards. (A) On each trial, head-fixed mice turn a wheel with their forepaws to move a visual stimulus to the center screen. The stimulus appears randomly on each trial on the left or right, at random contrast. (B) Single-trial timeline. (C) The available reward size depends on the stimulus in a manner that alternates over blocks, where correct left choices are rewarded more than right ones (left blocks, *green matrix*) or vice versa (right blocks, *orange matrix*). (D) Psychometric performance for left blocks (green) and right blocks (orange) for six mice over 80 sessions (46,236 trials), plotted as the proportion of right choices for each contrast condition. Mice choose according to the stimulus location and contrast, but are also influenced by the block: they choose left more often in left blocks, and choose right more often in right blocks. Thin lines, mean performance for individual mice; bold lines and error bars, mean ± s.e. across mice. (E) The difference in performance between left and right blocks for the same mice and sessions shown in (D). Mice are more influenced by the block structure (larger right–left difference) on low-contrast trials than high-contrast trials. Thin lines, mean delta performance for individual mice; bold lines and error bars, mean ± s.e. across mice.

Mice mastered this task after several weeks of training (43 ± 7 days between initial water restriction and first imaging session, mean ± s.e., *n* = 6 mice), efficiently combining current sensory evidence with expected reward (Figure 1D). In this task, high-contrast stimuli are unambiguous and should always be chosen, because choosing the other side gives no reward. Conversely, on low-contrast trials, where the stimulus side is ambiguous, mice should favor the side assigned by the block, as this leads to higher expected reward (Whiteley and Sahani, 2008; Lak et al., 2020). Consistent with this prediction, psychometric curves show larger behavioral effects of the block side on low-contrast trials than high-contrast trials (Figure 1E).

### MOs activity correlates with multiple task features

Neurons in frontal area MOs showed diverse and coordinated activity during task performance. We used multiplane two-photon calcium imaging in CaMK2a-tTA;tetO-GCaMP6s mice (*n* = 6) performing the task to record the activity (deconvolved fluorescence) of neurons in layer 2/3 of either the left or right hemisphere of MOs, across various fields of view (1.21 ± 0.08 mm anterior and 0.81 ± 0.02 mm lateral to bregma; mean ± s.e. *n* = 12) in a 4 mm cranial window centered at +1 mm anterior to bregma (Figure 2A). To visualize population activity in an unsupervised manner, we used rasterMap (Stringer et al., 2019), which arranges neurons so that those with correlated activity are placed close together. Analysis of individual trials revealed populations of neurons active in response to task events (Figure 2B). Plotting the mean peristimulus time histogram (PSTH) of all neurons revealed a striking structure, with neurons diversely activated or suppressed at different moments within the trial (e.g., pre-trial period, delay period, or response/feedback period, Figure 2C).

**Figure 2.**
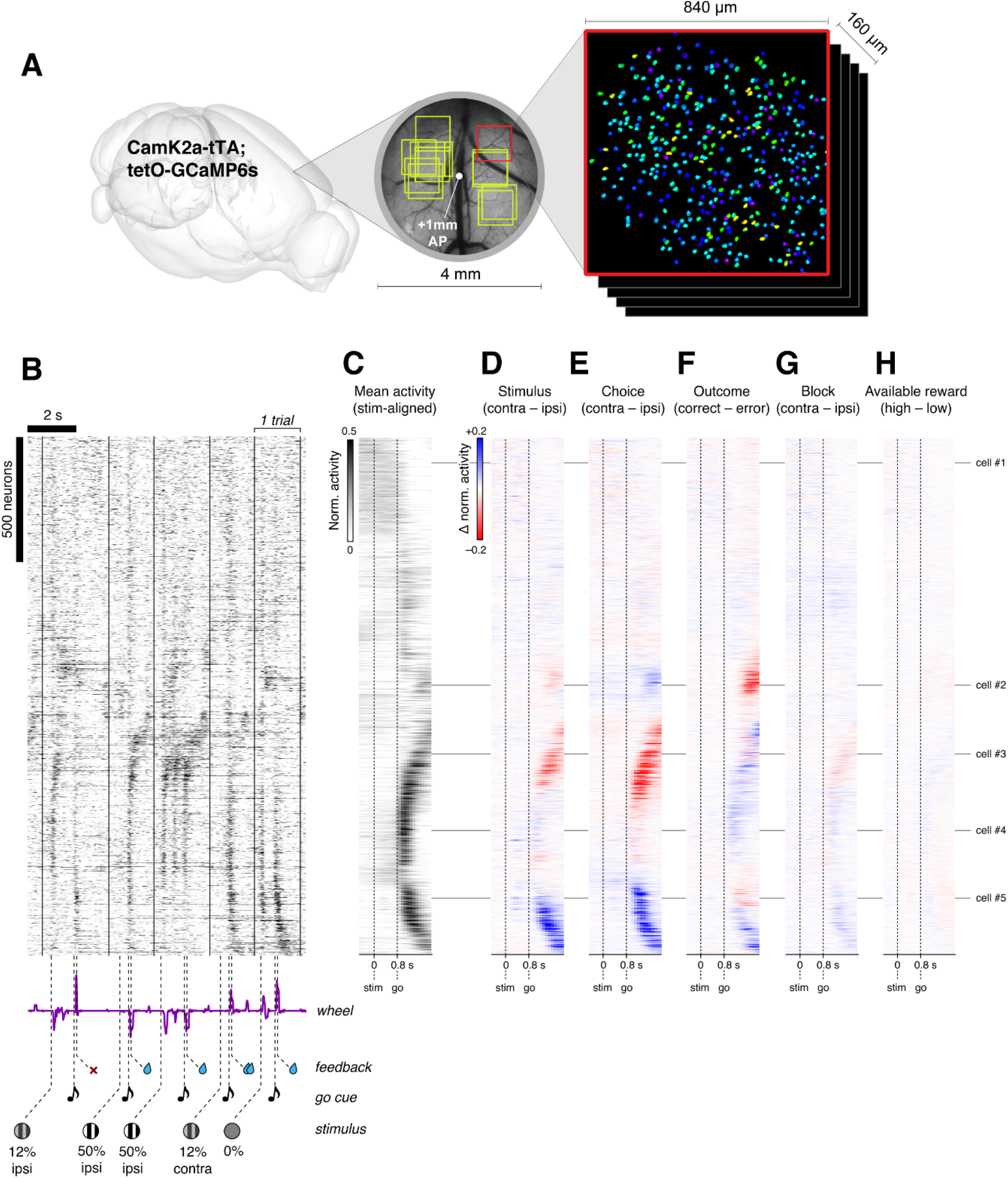
MOs activity correlates with multiple task features. (A) A 4mm round cranial window was implanted over MOs, and each imaging session targeted a ∼840 × 840 um field-of-view (FOV) within this window (yellow squares). The stereotaxic locations of twelve unique FOVs from six mice are shown relative to +1 mm AP in the cranial window (example brain image for context). Image registration, cell detection and fluorescence deconvolution were done offline using Suite2P (Pachitariu et al., 2016). (B) Raster plot showing activity of 1,957 neurons from an example recording session over five trials of the task. Neurons were organized vertically using rasterMap (Stringer et al., 2019) to place correlated neurons together. Solid lines denote individual trial starts, and task events are labeled along with the corresponding segment of wheel movement velocity. (C) Raster showing stimulus-aligned mean activity of the same population, sorted vertically as in (B), averaged over all trials from the session. (D) Raster showing the stimulus-aligned average difference between contra and ipsi stimulus trials. (E–H) Same as in (D), for the stimulus-aligned average difference between contra and ipsi choice (E), correct and error outcome (F), contra and ipsi reward block (G), and high- and low-volume available reward trials (H). Horizontal lines mark 5 example neurons analyzed in Figure 3.

As a first analysis of neural coding of task variables, we visualized the difference in mean response between trial types. This analysis revealed strong apparent correlates of population activity with stimulus side (contra vs. ipsi, Figure 2D), choice side (contra vs. ipsi, Figure 2E), and outcome type (correct vs. error, Figure 2F). By contrast, there appeared to be weaker, if any, neural correlates of reward block (contra vs. ipsi, Figure 2G) and available reward (high vs. low: the reward that would be given for a correct choice, depending on the combination of stimulus side and block but independent of the mouse’s actual choice, Figure 2H). However, because the task variables are themselves correlated, further analyses are required to isolate which cells’ activity relate to which task variables.

### MOs neurons encode stimuli, choices, and outcomes

To isolate which task features were encoded by individual neurons, we first used a kernel-fitting approach (Park et al., 2014; Steinmetz et al., 2019) (Figure 3A). We fit the activity of each neuron as a sum of eleven kernel functions time-locked to task events. Five of these kernels captured variations in amplitude and timing of visual responses (one kernel for high- or low-contrast stimuli on each side and one for zero-contrast trials), and were locked to stimulus onset (−0.5– 2s relative to stimulus onset). A “block” kernel and an “available reward” kernel captured possible differences between contra and ipsi blocks, and between trials of low vs. high available reward, at all moments within the trial (−0.5–2s relative to stimulus onset). Two additional kernels captured correlates of left and right movements (−0.5–1s relative to movement onset): an “action” kernel triggered by a movement in either direction, and a “choice” kernel capturing differences in activity between left and right movements (Steinmetz et al., 2019). Finally, two “outcome” kernels (−0.5–1s relative to feedback time) captured correlates of feedback delivery/reward consumption on correct and error trials. Comparing the model’s predictions with observed firing showed that these kernels were sufficient to capture the activity of MOs neurons (Figure 3A).

**Figure 3.**
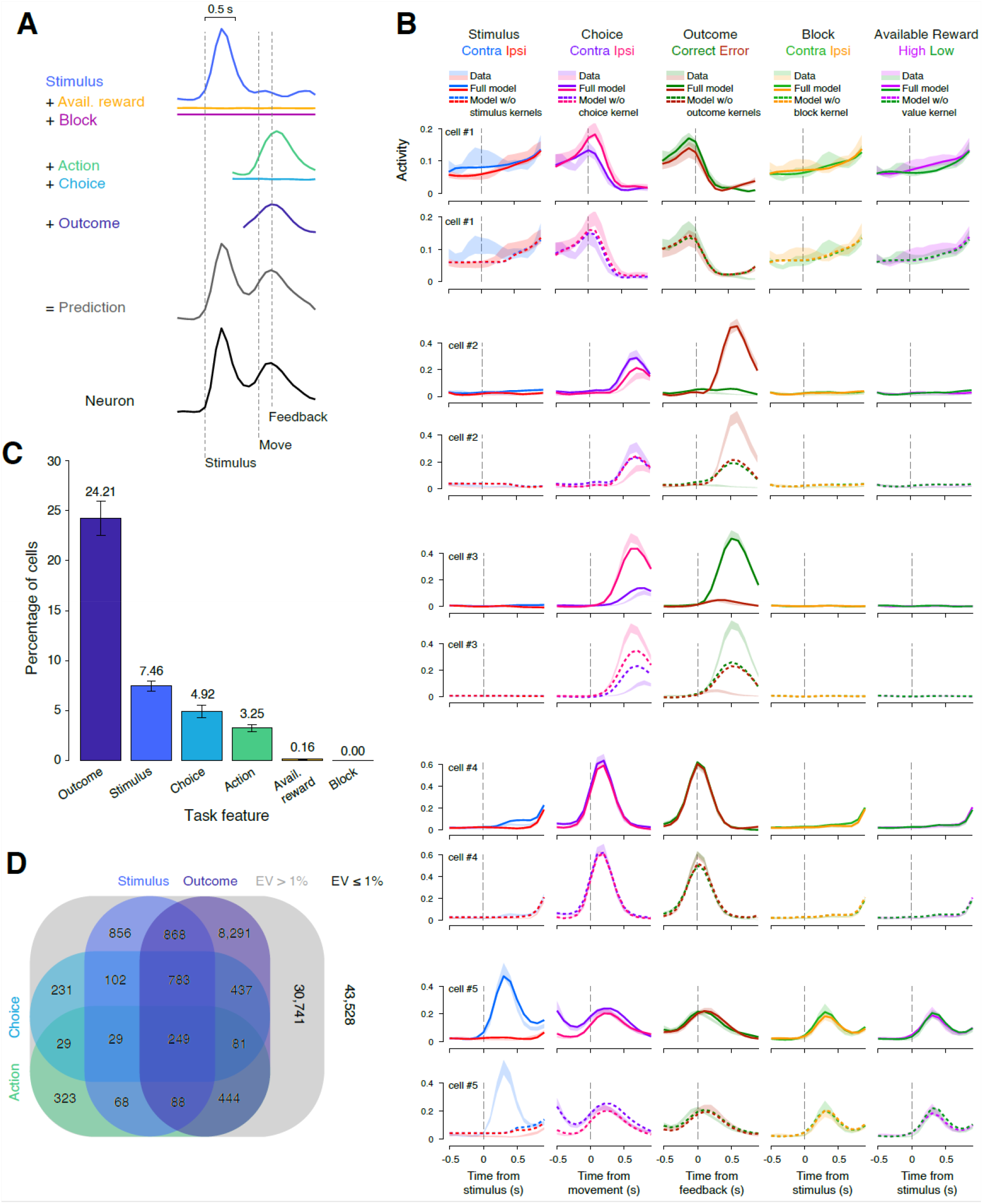
MOs neurons encode stimuli, choices, and outcomes. (A) Activity on each trial is modeled as a sum of cell-dependent kernels aligned to the relevant event onset time. Black trace: firing rate for an example cell averaged over trials. Colors: individual kernels averaged over trials. Gray: prediction from kernel sums. (B) Example nested fitting analysis for the five example neurons labeled in Figure 2. Firing rate was averaged across the trial types indicated (shaded regions, mean ± s.e. across trials): contralateral or ipsilateral stimuli (blue or red), contralateral or ipsilateral choices (purple or pink), correct or error outcomes (green or brown), contra or ipsi block (green or orange), and high or low available reward (magenta or green). Solid lines: cross-validated prediction using all kernels. Dashed lines: predictions refit from a model excluding the indicated kernel. Note for each of these neurons the good fit of the full model is lost when excluding one of the kernels (stimulus, choice and/or outcome), indicating that these neurons have specific stimulus-, choice-, and/or outcome-related activity that cannot be explained by the remaining kernels. (C) Percentage of cells (mean ± s.e.) across 36 sessions (*n* = 89,092) in which >1% of variance is explained by the given task event kernel, assessed through nested fitting analysis. Neurons with outcome kernels dominate the population, followed by stimulus, choice, and action. Very few available reward or block neurons were identified. (D) Neurons often encode more than one task feature, as shown by overlapping regions of the Venn diagram. A 4-way Venn diagram was used for simplicity due to the very small numbers of neurons encoding available reward or block.

To determine which task features were needed to predict the activity of which individual neurons, we used a nested fitting procedure (Figure 3B). This approach asks whether, for each cell and each feature, prediction of the cell’s activity using all feature kernels is more accurate than prediction using all but one feature kernel to be tested. We deemed that a feature was necessary to predict a cell’s activity if it increased explained variance by >1% on held-out data. For example, in the neurons labeled in Figure 2, the full kernel model successfully captured a range of different firing dynamics across neurons, and the nested fitting protocol successfully quantified the contributions of various task features to individual cells (Figure 3B). The fraction of cross-validated variance explained by the kernels was frequently small, even for neurons whose mean rates they accurately predicted (e.g., 2.2%, 17.5%, 45.1%, 41.7%, and 28.7% for Cells 1–5, Figure 3B), as expected from trial-to-trial variability due to factors such as the encoding of task-independent variables (Musall et al., 2019; Stringer et al., 2019).

Accounting for the activity of MOs neurons typically required kernels for stimulus, choice, action, and outcome, but rarely required kernels for available reward and reward block. The most commonly required kernels were those for outcome (24.2 ± 1.7% of neurons in a session, mean ± s.e., *n* = 36 sessions), followed by stimulus (7.5 ± 0.5%), choice (4.9 ± 0.6%), and action (3.3 ± 0.3%). Very few neurons required the kernels for available reward (0.16 ± 0.05%) or reward block (0.002 ± 0.002%) (Figure 3C). Neurons often encoded more than one task feature (Figure 3D), reflecting a mixed-selectivity code (Rigotti et al., 2013).

### MOs significantly encodes all task variables except reward block

The kernel analysis estimates how many neurons encode specific task features but not how well the population encodes those features. Furthermore, it does not provide a rigorous measure of statistical significance. Such a measure should take into account potentially spurious correlations arising from slow drifts in neural activity and unrelated slow drifts in behavioral timeseries (Harris, 2020a).

We solved this problem by decoding task variables from neural activity and comparing decoder performance to surrogate data where there was no relationship between neural and task variables (Figure 4). We trained L1 regularized logistic regression classifiers to predict binary task variables on each trial: stimulus side (contra vs. ipsi), choice side (contra vs. ipsi), outcome (correct vs. error), block (contra vs. ipsi) and available reward (high- vs. low-volume). The classifiers operated on population neural activity at a range of time points across the trial, aligned to cue onset (−0.5–3s). With this approach, we were able to determine when, over the course of a trial, a task feature could be decoded from the neural population. We quantified the performance of logistic decoders as the cross-validated log_2_ likelihood ratio of the “full” decoder to a “naïve” decoder without access to neural activity (which predicted the probability of each task feature simply by its frequency in the training set). This predictability measure provides an estimate of the mutual information between neural population activity and the task variable (Kjaer et al., 1994; Harris et al., 2003). The predictor yielded positive predictability values for all variables following the Go cue, and for choice and stimulus side during the delay period (Figure 4A-E, black). To test whether these predictions were significantly larger than would be expected by chance, we compared predictions to surrogate data, using one of two methods according to whether the task variable being predicted was randomized.

**Figure 4.**
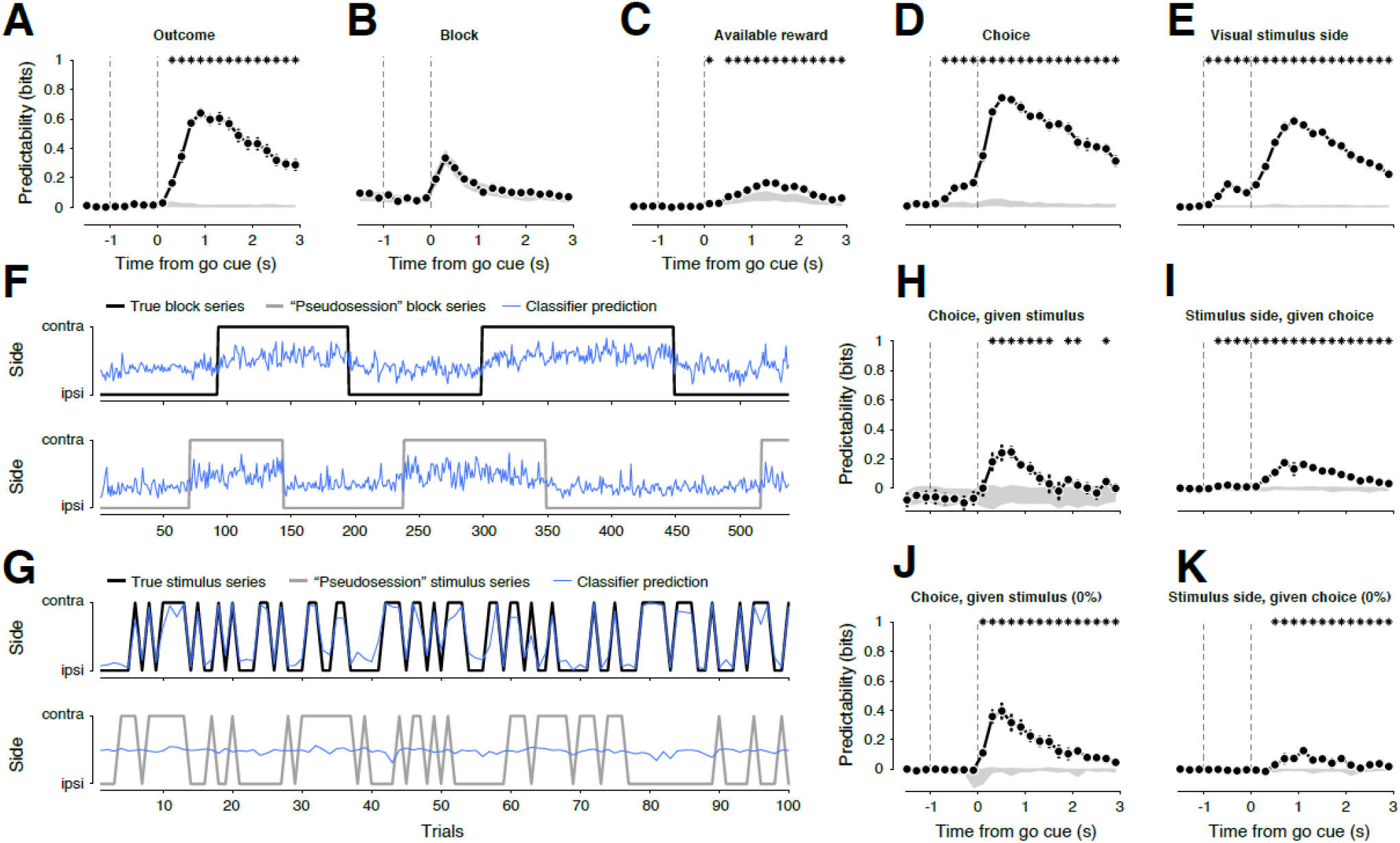
MOs significantly encodes all task variables except reward block. (A–E) Predictability of task variables from neural activity at various timepoints in the trial. Shown are predictability of outcome (A), reward block (B), available reward (C), choice (D) and stimulus side (E). Black data points, mean ± s.e. of predictability for actual neural and task data (*n* = 36 sessions); gray filled region, 2.5^th^–97.5^th^ percentile of null distribution generated by predicting “pseudosessions” where data were randomly regenerated (B,C,E) or linearly shifted in time (A,D). Asterisks denote timepoints when the actual task variable is predicted significantly better than the null ensemble. Vertical dashed lines denote earliest possible stimulus onset time (−1 s) and the time of Go cue (0 s). (F) Top: true block sequence (black line) and its prediction from neural activity (blue curve), for each trial of an example session. Bottom: randomly regenerated pseudoblock sequence (gray line) and its prediction from neural activity (blue curve). (G) Same as (F) but for stimulus sequence in the same session. Only the first 100 trials are plotted. (H) Predictability of choice side from neural activity, taking into account the common effect of stimulus side, plotted as in (A-E). (I) Predictability of stimulus side from activity, taking into account the common effect of choice side. (J–K) The same decoder as in (H–I), but this time trained on non-zero contrast trials and validated on held-out zero-contrast trials.

We first tested the neural decoding of randomized task variables, and found significant decoding of stimulus, and available reward but not of reward block. In our task, reward block, stimulus side, and available reward size are randomly generated independent of the subject’s choices. To test the significance of decoding these variables, we used a pseudosession test (Harris, 2020a), which compares the decodability of the actual task variable series against a null ensemble obtained by repeatedly predicting the task variable series that would have occurred in “pseudosessions” generated using the same probability rules (Figure 4B,C,E, gray). For block coding, the apparent predictability of the block side from neural activity was not significant: the measured neural activity could be used to predict randomly-generated pseudoblocks just as well that it could be used to predict the block sequence that the mouse experienced (Figure 4B, F). For both the real block sequence and the pseudoblock sequences, predictions were strongest from neural activity immediately after the Go cue, presumably because activity was highest at this time (Figure 2C). Even at this time, however, the real block sequence was no better predicted than the ensemble of pseudoblock sequences, indicating that neural activity was no more correlated with the block sequence than expected by chance. By contrast, prediction of the true stimulus side sequence far exceeded prediction of a pseudosession sequence (Figure 4E, G). Prediction of available reward size was weak but significant at times after the Go cue, when rewards were being consumed (Figure 4C). This coding of available reward size thus most likely reflects an effect of the physical volume of reward delivered, rather than a cognitive coding of expected reward, which would also occur during the delay period.

We then considered task variables that depend on the animal’s behavior—choice and outcome—and found coding of choice starting in the delay period, and of outcome starting from the Go cue (which, due to the slow sampling of calcium imaging, cannot be distinguished from the movement and reward times). To test for significant decoding of these variables that are not randomized but depend on the animal’s choices, we used a linear shift method, which compares the predictability of the genuine variable to a null ensemble generated by shifting neural activity forward or backward between trials (Harris, 2020b). This test revealed strong coding of both outcome and choice following Go cue onset, with choice also more weakly coded during the delay period (Figure 4A, D).

However, the apparent relationship between delay-period neural activity and the upcoming choice could be explained by their common dependence on the visual stimulus. To test this, we considered two decoders: a ‘full’ decoder that predicted one task variable from the other plus neural activity (e.g., predicting choice from stimulus plus neural activity) and a ‘naïve’ decoder that predicted one task variable from the other (e.g., predicting choice from stimulus). We then computed the partial predictability of one task variable given the other by subtracting the log_2_ likelihoods of the two predictions. This analysis confirmed the predictability of stimulus and choice after the Go cue (Figure 4H,I), but found no significant encoding of choice in the earlier delay period (Figure 4H). This result suggests that during the delay period, the predictability of choice from MOs activity alone (Figure 4D) could be explained by the predictability of choice from stimulus. To further investigate this possibility, we restricted the previous analysis to trials of zero contrast, when there was no visible stimulus at all. In these trials, the “stimulus side” variable—which indicates which choice would be rewarded—is invisible to the mouse. Confirming our previous results, we found that choice was significantly coded only after the Go cue, and not during the delay period (Figure 4J). As for the stimulus side, as expected, it was not encoded during the delay period (the stimulus was invisible in these trials), but intriguingly, it was encoded following the Go cue (Figure 4K). As we show next, this encoding could be explained by a nonlinear combination of choice and outcome signals.

### MOs encodes exclusive-or (XOR) of choices and outcomes

At zero contrast, when the stimulus is invisible, it is possible to infer the stimulus-side variable from the animal’s choice and the trial outcome via an exclusive- or (XOR) operation: a rewarded left choice or unrewarded right choice implies the stimulus side was on the left, while a rewarded right choice or unrewarded left choice implies right. It is well known that XOR operations cannot be performed by linear readout (Minsky and Papert, 1969), yet we found it possible to linearly predict stimulus side from MOs activity even on trials of zero contrast (Figure 4K). This therefore supports the idea that the MOs population encodes a nonlinear interaction between the choice and outcome variables. Examining neural activity rasters of the four possible choice-outcome combinations revealed example MOs neurons that appeared to encode choice-outcome interactions on zero-contrast trials (Figure 5A–D). To confirm this statistically, we deployed a two-way ANOVA test that included a main effect term for choice and outcome independently, plus an interaction term. Of the entire population of 89,092 neurons, 8,919 cells had a significant main effect of outcome, 8,105 had a significant main effect of choice, and 4,912 significantly coded an interaction of choice × outcome (*p* < 0.05, F-test; Figure 5E). To visualize the geometry of coding for the choice, outcome, and their interaction (XOR), we trained logistic decoders to predict these three variables on each trial, training each on all non-zero-contrast trials and validating on held-out zero-contrast trials. When labeled by trial type, the points formed distinct corners of a 3-dimensional tetrahedron (Figure 5F), unlike the planar geometry that would reflect a linear code (Bernardi et al., 2020). All three decoders performed well above chance, as assessed against pseudo-decoders that were trained to predict trial identities using trial-shifted neural data (Figure 5G).

**Figure 5.**
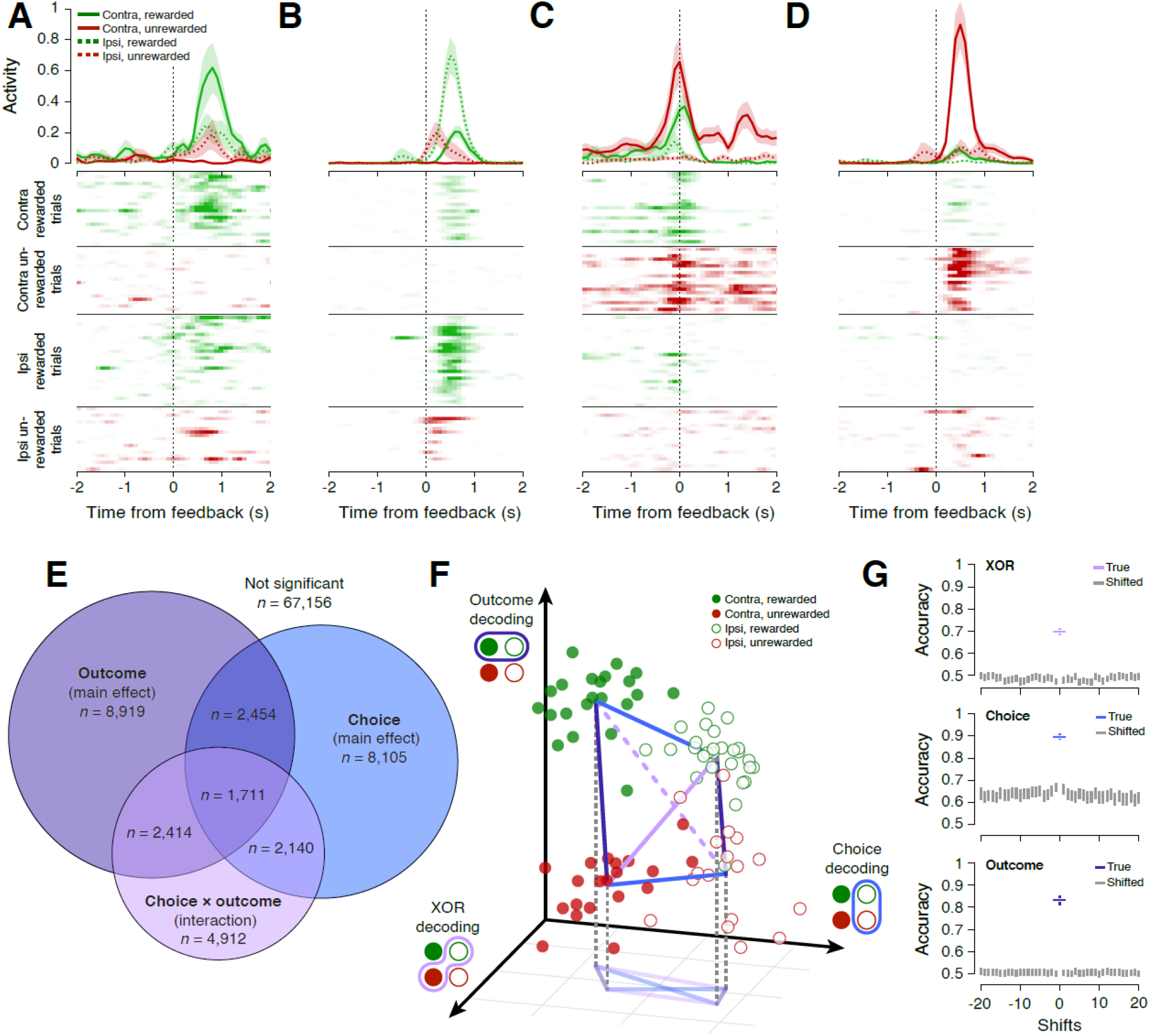
MOs encodes exclusive-or (XOR) of choices and outcomes. (A–D) Peri-event time histograms and rasters of activity time-aligned to feedback onset, shown for four example neurons, split by the four possible conditions on zero-contrast trials: rewarded contra choices, unrewarded contra choices, rewarded ipsi choices, unrewarded ipsi choices. Neurons do not linearly code choice and outcome, as indicated by different responses to each condition. (E) Two-way ANOVA for all neurons (*n* = 89,092) reveals cells with significant main effects for choice and outcome, but also neurons with a significant interaction effect, which can be used to solve the XOR function. The inferred stimulus location (i.e., the side to which a choice would have been rewarded) is given by the XOR of choice and outcome, even on zero-contrast trials. (F) Decoding of outcome, choice, and inferred stimulus location with one point for each zero-contrast trial (x, y, and z axes, respectively) for an example session. If XOR decoding were not present, the tetrahedron would be two-dimensional. (G) Decoder accuracy for XOR (top), choice (middle), and outcome (bottom) across all sessions. Gray vertical bars denote the mean ± s.e. across mice for the accuracy of pseudo models trained on a series of trial-shifted data (shifts = -20 to +20 trials); the colored cross is the mean ± s.e. across mice for the accuracy of the true model trained on non-shifted data (shift = 0).

## Discussion

We found that in a visual decision-making task where reward sizes varied in blocks, the MOs population encoded sensory stimulus, available reward size, choice direction, and trial outcome (correct vs. error), However, the encoding of upcoming choice direction in the delay period between stimulus presentation and the Go cue could be explained by encoding of the stimulus side. Encoding of available reward size was weakly present during the feedback period when rewards were consumed. Encoding of the reward block side was not statistically significant.

Data from experiments such as this one require careful statistical analysis to avoid detecting “nonsense correlations” that can occur with methods assuming statistical independence between trials (Yule, 1926; Elber-Dorozko and Loewenstein, 2018; Harris, 2020a). For example, predicting the block variable from population activity following the Go cue on each trial yielded an apparent correlation, even using cross-validation to assess fit quality. However, this apparent prediction was not genuine. Cross-validation is not sufficient to eliminate predictions between timeseries that show independent slow changes (Harris, 2020a); indeed, it was possible to predict a fake sequence of block variables, drawn randomly using the same probability rules as the original blocks (a “pseudosession”) just as accurately as the block sequence the mouse experienced. This shows that the apparent predictability of the block variable is no more than what would be expected due to random coincidence between slowly drifting neural activity and temporally extended block variables.

During the delay period between visual stimulus and motor response, neural activity in MOs correlated with both the stimulus side and the mouse’s upcoming choices, but this could be explained by encoding of only the visual stimulus. Stimulus and choice are themselves correlated, so delay period coding could in principle reflect neural correlates of the visual stimulus (which remains on the screen), the mouse’s choice, or preparatory activity for a movement to be released when the Go cue sounds. Our data suggest that a neural correlate of the visual stimulus is sufficient to explain activity in the delay period: if delay-period activity encoded the upcoming choice then activity should predict upcoming choice even on zero-contrast trials, but this was not the case. This finding differs from previous findings in memory tasks, in which delay-period activity correlates better with upcoming movement than sensory stimuli (Erlich et al., 2011; Li et al., 2016).

We hypothesize this difference from previous results arises from differences in task design. Our task is not a memory task: the stimulus remains on the screen throughout the delay period, so the mouse does not need to commit to a choice until the Go cue sounds. In memory tasks, where the stimulus is not physically present during the delay period, the subject must either memorize the stimulus location or commit to an action during the delay period. Studies of memory tasks have distinguished these two possibilities by examining error trials, or by using tasks in which auxiliary stimuli direct the subject to respond differently to the same cue, with mixed results. For example, in a task in where the shape of the fixation spot indicated whether monkeys should saccade toward or away from a visual cue (Funahashi et al., 1993), most frontal neurons encoded the cue location with a minority encoding the upcoming movement direction. In a memory-guided orienting task for rats, analysis of error trials suggested that most neurons in frontal orienting fields predicted upcoming motion (Erlich et al., 2011), and a similar result was found in a left/right licking task in head-fixed mice (Li et al., 2016). In contrast, in a task requiring monkeys to compare an initial sensory stimulus to a second stimulus following a delay, frontal cortical activity during the delay period correlated with the initial stimulus (Romo et al., 1999).

The lack of significant coding of reward block side may also be specific to this task. In our task, block side controls the amount of reward given for a correct choice, but does not affect the probability of a stimulus appearing on either side: the mouse is always rewarded for choosing the side with the visual stimulus, and never for the side without it. Psychometric performance shows that stimulus strongly influences animal choices and block only weakly, so one possibility is that the encoding of reward block is smaller than we could detect in our analyses. Another possibility is that because the stimulus remains onscreen throughout the trial, the animal’s strategy is different and does not require MOs to encode this variable at all.

Indeed, in contrast to our results, a maze-based two-armed bandit task showed MOs correlates of choice (and thus likely of reward block) prior to the choice itself (Sul et al., 2011). Furthermore, data from our lab using a head-fixed two-armed bandit task—with the same wheel behavior as the present task but no visual stimuli—does show significant coding of block side during the intertrial interval (Lebedeva et al., 2022). We suggest that in two-armed bandit tasks such as these, committing to a particular choice in advance of choice execution is an optimal strategy; therefore delay-period MOs activity will correlate with choice and block in bandit tasks, but not in the current task. This supports the hypothesis that the role of frontal cortex in different tasks is flexible and can differ depending on the behavioral strategy used for each task (Duncan, 2001).

In summary, we observed diverse neural correlates in MOs, including trial outcome, action, choice, visual stimulus, and even a nonlinearly coded inference of stimulus side from outcome and choice on trials with no visual stimulus. The correlates of MOs activity we found in this task differ from correlates observed in other choice tasks, suggesting that frontal cortex can be rewired in a manner appropriate to the behavioral demands of each task. Understanding the precise neural correlates found in different tasks will be essential towards deciphering the learning rule that frontal cortex uses to form these representations.

## Acknowledgments

Many thanks to Laura Funnell and Yeqing Wang for animal training, to Bex Terry and Anne Ritoux for data preprocessing, and to Anna Lebedeva, Kevin Miller, and Max Shinn for discussions. This research was funded by Marie Skłodowska-Curie IF No. 795846 (to L.E.W.), Wellcome Trust Award No. 213465 (to A.L.), Wellcome Trust Award No. 205093 (to M.C. and K.D.H.), and ERC advanced award No. 694401 (to K.D.H.). M.C. holds the GlaxoSmithKline / Fight for Sight Chair in Visual Neuroscience.

## Contribution statement

Contributor roles taxonomy from https://credit.niso.org

|  | LEW | AL | MC | KDH |
| --- | --- | --- | --- | --- |
| **Conceptualization** | ● | ● | ● | ● |
| **Data curation** | ● |  |  | ● |
| **Formal analysis** | ● |  |  | ● |
| **Funding acquisition** | ● |  | ● | ● |
| **Investigation** | ● |  |  |  |
| **Methodology** | ● |  |  | ● |
| **Project administration** | ● |  |  | ● |
| **Resources** |  | ● |  |  |
| **Software** | ● |  |  |  |
| **Supervision** |  |  | ● | ● |
| **Visualization** | ● |  |  |  |
| **Writing – original draft** | ● |  |  | ● |
| **Writing – review and editing** | ● | ● | ● | ● |

## Data and code availability

Data and code are available at https://figshare.com/projects/Mouse_frontal_cortex_nonlinearly_encodes_sensory_choice_and_outcome_signals/166622.

## Methods

### Experimental procedures

All experiments were conducted according to the UK Animals Scientific Procedures Act (1986) under appropriate project and personal licenses.

### Surgeries

Six transgenic adult mice (60 days or older; 4 male) expressing GCaMP6s in excitatory neurons (CaMK2a-tTA;tetO-GCaMP6s) were implanted with a custom metal head plate with 7mm circular well, inset with a 4mm glass cranial window set over secondary motor area (MOs). Animals were first anesthetized with isoflurane and an ophthalmic ointment was applied to the eyes, and injections of carprofen and dexamethasone were administered. The hair on the head at the planned incision site was shaved away, and the mouse was transferred to a stereotaxic apparatus where its skull was secured with ear bars, and body kept on a feedback-controlled heating pad (ATC2000, World Precision Instruments, Inc.). The scalp was cleaned with 70% ethanol to remove loose hairs and other detritus, after which a lidocaine ointment was applied. Following a final application of iodine and ethanol, the scalp was excised, and the edges of the incision were sealed to the skull with a cyanoacrylate adhesive. The head plate was attached medially to the skull at +1 mm bregma using dental acrylic resin. A craniotomy was made using a 4mm biopsy punch (Integra, Miltex) and angled probe (10032-13, Fine Science Tools) to remove the medial bone flap, taking care not to disturb the central blood vessel or the surrounding dura. A double coverslip (4mm inner, 5mm outer) was placed into the craniotomy, held in place with a toothpick and bonded to the skull with cyanoacrylate adhesive. Finally, all exposed skull was covered with additional dental cement to seal the cranial window and secure the head plate. Post-operative pain prevention was maintained with carprofen in drinking water on the three following days.

### Behavioral task

Behavioral training started at least 7 days after the head plate implantation surgery. Animals were handled and acclimatized to head fixation for 3 days, then began water restriction and training in a 2-alternative forced choice visual detection task (Burgess et al., 2017), after which they were introduced to unequal water rewards for left or right choices (Lak et al., 2020). Three computer screens surrounded the mouse, spanning -135 to +135 v° along the azimuth axis and -35 to +35 v° along the elevation axis. After a pre-trial period of 0.5–1 s where the mouse must keep the wheel still, a vertical grating stimulus (square wave with Gaussian window) of varying contrast (100, 50, 12, 5, 0%) appeared on either the left or right monitor with 50% probability (0% contrast trials were also assigned to the left or right screen with the same probability). After a delay period of 0.8–1.0 s, in which the mouse must continue to keep the wheel still, a brief Go cue played (0.1 s, 8 kHz) after which the mouse could report its decision by turning the wheel located underneath its forepaws. Impulsive movements, in which the wheel was moved before the Go cue, caused the delay period to reset until quiescence was achieved. Wheel movements after the Go cue drove the stimulus across the monitor(s). A reward was delivered if the stimulus reached the center of the middle monitor (a correct trial) followed by a 1 s post-feedback period during which the stimulus remained onscreen for 0.5 s. By contrast, a 0.5 s white noise was played if the stimulus was moved offscreen in the opposite direction (an error trial), followed by a 2 s post-feedback period. All trials had a further 1 s inter-trial interval after the feedback period. Unequal rewards, in which either left or right correct choices elicited larger rewards (2.4 µL versus 1.2 µL of water), occurred in blocks of 125–225 trials (drawn from a uniform distribution). Only those trials without impulsive movements were included in subsequent analyses.

The behavioral experiments were controlled by custom-made software written in MATLAB (Mathworks) which is freely available (Bhagat et al., 2020). Instructions for hardware assembly are also freely available (https://www.ucl.ac.uk/cortexlab/tools/wheel).

### Two-photon imaging

Calcium imaging was performed in trained animals while they were head-fixed and performed the behavioral task. Layer 2/3 in MOs was imaged using a commercial two-photon microscope (Bergamo II, Thorlabs Inc) controlled by ScanImage. A Ti:sapphire laser (Chameleon Vision, Coherent) was set to a 920 nm wavelength, and the beam was focused with a 16x water-immersion objective (0.8 NA, Nikon). Images were acquired at a frequency of 30 Hz across six planes (5 Hz per plane), a resolution of 512 × 512 pixels, with a frame width of 840 µm. The fly-back plane was excluded from further analysis.

### Cell detection

Registration, cell detection, neuropil correction, and deconvolution of the two-photon imaging data were carried out using Suite2P (Pachitariu et al., 2016). Imaged planes were aligned with non-rigid registration (four blocks, 128 × 128), and spiking activity was deconvolved from calcium fluorescence using a kernel with a timescale of 2 s. Deconvolved spike traces were used for all subsequent analyses.

### Kernel regression analysis

To identify which task features were encoded by which neurons, we began by fitting a kernel-regression model. In this analysis, the firing rate of each neuron is modeled as a linear sum of kernels aligned to task events. For the current study, we used kernels for reward block, available reward, stimulus onset, choice, action onset, and outcome.

For each kernel to be fit on a task event, *d*, we constructed a Toeplitz predictor matrix *P*_*d*_ of size *T* × *L*_*d*_, in which *T* is the total number of time points in the training set, and *L*_*d*_ is the number of lags required for the event kernel. This matrix contains diagonal stripes where *P*_*d*_(*t,i*) = 1 if *t* – *i* ∈ *d* and 0 otherwise. Predictor matrices were defined similarly for all other kernels, then all predictors were horizontally concatenated to yield a global prediction matrix *P* of size *T* × 235.

To regularize the kernel fit, we first applied reduced-rank regression (Steinmetz et al., 2019), to factorize the kernel matrix *K* into the product of a 235 × *R* matrix *B* and a *R* × *N* matrix *W*, minimizing the total error: *E* = ‖*F* – *PBW*‖^2^. The *T* × *R* matrix *PB* may be considered as a set of temporal basis functions, which can be linearly combined to estimate each neuron’s firing rate. Reduced-rank regression ensures that these basis functions are ordered, so that predicting population activity from only the first *r* columns will result in the best possible prediction from any rank *r* matrix.

To derive neuron *n*’s kernel functions, we estimated a weight vector *w*_*n*_ to minimize *E*_*n*_ = |*f*_*n*_ – *PBw*_*n*_|^2,^ where *f*_*n*_ is neuron *n*’s firing rate, *PB* is the set of basis functions, and *E*_*n*_ is the error for neuron *n*. We used elastic-net regularization using the package glmnet (Friedman et al., 2010) (compiled for MATLAB 2020b; https://github.com/lachioma/glmnet_matlab) with parameter *α* = 0.5, determining the optimal number of columns *r*_*n*_ of *PB* to keep when predicting neuron *n* as the one giving optimal performance on five-fold cross validation. The kernel functions for neuron *n* were then unpacked from the 235-dimensional vector obtained by multiplying the first *r*_*n*_ columns of *B* by *w*_*n*_. Neurons with total cross-validated variance explained of <1% were not considered responsive.

To assess the selectivity of individual neurons for each kernel, we used a nested approach as in (Steinmetz et al., 2019). We first fit the activity of each neuron, excluding the kernel to be tested, using the reduced-rank regression procedure above (including deriving a new basis set). We subtracted this prediction from the raw firing rate to obtain the residuals, representing aspects of the neuron’s activity not explainable by the other kernels. We then repeated the reduced-rank regression procedure but this time using the residual firing rates as the independent variable, and using only the test kernel. The cross-validated quality of this fit determined the variance explainable only by the test kernel. If this variance explained was >1%, the neuron was deemed selective for that kernel.

### Decoding task variables from the neural population

We trained L1 (lasso) regularized logistic regression models to predict trial-by-trial task features (stimulus on the contralateral or ipsilateral screen, contra or ipsi choice by the mouse, correct or error outcome, contra or ipsi block, or high- or low-volume reward) from population neural activity.

To predict a binomial task feature (represented as 1 or 0) at time t, we trained a logistic regression model that included as predictors a matrix of neural activity at time *t* (nTrials x nNeurons). We used MATLAB cvglmnet (three-fold cross-validation), using the lambda giving optimal cross-validated performance. We compared this full model to a naïve model whose trial-by-trial prediction included no neural data and predicted the probability of each task feature simply by its frequency in the training set. The cross-validated log_2_ likelihood ratio of these models measures the predictability of the task feature from neural activity, and provides a lower-bound estimate of the mutual information between them (Harris et al., 2003; Itskov et al., 2007).

To separate the neural correlates of stimulus location and choice, which are correlated in the task, we additionally utilized a more rigorous naïve model for predicting stimulus side or choice direction that accounts for this correlation. For these models, we trained a full model that included as predictors a matrix of neural activity at time *t* (nTrials x nNeurons) and a behavioral matrix (for stimulus side, this was a nTrials x 1 vector of the animal’s choices; for choice direction, this was a nTrials x 8 “one-hot encoding” design matrix of stimulus values). We used MATLAB cvglmnet to automate lambda selection for regularization of predictors in the neural matrix, but excluded regularization of the behavioral predictor(s) (“options.exclude” and “options.penalty_factor”). We compared this full model to a naïve, unregularized model using the behavioral predictor(s) only, reporting the difference to quantify how much neural activity improved predictions.

To examine predictions of stimulus side and choice on zero-contrast trials, we designed neural and behavioral predictor(s) as above, but split data into custom training and validation sets, where the validation set contained all zero-contrast trials(∼20%), and the training set was all other trials (∼80%).

Predictions of choice, and outcome using neural data were tested for significance using a linear-shift method, which misaligns neural data from trial data in a series of increasing backward or forward trial shifts (Harris, 2020b). Predictions were considered significant if the predictability using trial-aligned neural data was larger than the predictability computed for all shifts of up to 20 trials in either direction, which provides a conservative test at *p* < 0.05 significance. Predictions of stimulus, block and available reward size were tested for significance by regenerating 100 pseudoseries of task events using the same statistics deployed during the task; significance was achieved if the predictability between full and naïve models using true task event series was >95% the predictability values computed from the 100 pseudoseries, which provides a conservative test at *p* < 0.05 significance.

### Decoding XOR (putative stimulus) from the neural population

To determine how well the neural population could determine the inferred stimulus side on zero-contrast trials, we trained three decoders to predict animal choice, trial outcome, and inferred stimulus location, which is an XOR operation based on the combination of choice and outcome. To predict choice direction (represented as +1 or -1), we trained a logistic regression model that included as predictors a matrix of mean neural activity in the 0-1 s period after feedback (nTrials x nNeurons). We used MATLAB cvglmnet to automate lambda selection for regularization. We split data into training and validation sets, where the validation set contained all trials of zero contrast (∼20% of trials), and the training set was all other trials (∼80% of trials). We used the three weighted sums of neural activity on each trial, prior to logistic transformation, to assign each trial to a point in 3-dimensional space (Figure 5F). The accuracy of each classifier was computed by comparing the sign of the true values of the task variable to the sign of the predictor. To test whether the prediction was statistically significant we used a linear-shift method (Harris, 2020b). Predictions were considered significant if the accuracy of the model trained on trial-aligned neural data was greater than the accuracy computed for all 40 models trained on trial-shifted neural data (shifts up to 20 in both directions), which provides a conservative test at *p* < 0.05 significance. This same process was repeated to predict trial outcome and inferred stimulus location.

